# Critical Cell Spacing Drives Phase Transition in Matrix-Mediated Tissue Condensation

**DOI:** 10.1101/2024.11.05.622090

**Authors:** Xiangjun Peng, Yuxuan Huang, Wenyu Kong, Yanan Du, Elliot L. Elson, Xi-Qiao Feng, Guy M. Genin

## Abstract

Biological tissues exhibit phase transitions governed by mechanical feedback between cells and their extracellular matrix (ECM). We demonstrate through bio-chemo-mechanical modeling that this emergent behavior arises from competing physical effects: increasing matrix stiffness enhances individual cell activation while simultaneously weakening long-range mechanical communication. This competition establishes a critical cell spacing threshold (80-160 *µ*m) that precisely matches experimental observations across diverse cell types and collagen densities. Our model reveals that the critical stretch ratio at which fibrous networks transition from compliant to strain-stiffening governs this threshold through the formation of tension bands between neighboring cells. These tension bands create a mechanical percolation network that drives the collective phase transition in tissue behavior. Our model explains how fibrous architecture controls emergent mechanical properties in biological systems and offers insight into both the physics of fiber-reinforced composite materials under active stress, and into potential mechanical interventions for fibrotic disorders.

---

Collective phase transitions emerge in many physical systems when local interactions give rise to global ordering. In biological tissues, a striking example occurs when cells embedded in a fibrous extracellular matrix (ECM) undergo a sharp transition from disordered to ordered states at critical cell densities^1–5^. This transition manifests as dramatic tissue condensation, where the material contracts to a fraction of its original volume when cell spacing falls below a critical threshold of 80-160 *µ*m^5^. Despite the fundamental importance of this phenomenon to tissue mechanics, the physical principles governing this sharp bifurcation remain poorly understood.

The mechanical interplay between cells and their surrounding fibrous network presents a fundamental paradox: while individual cellular contractile responses increase monotonically with substrate stiffness, collective behaviors exhibit sharp transitions characteristic of critical phenomena. Recent studies have established the concept of optimal substrate stiffness for processes such as cell adhesion and migration^6–8^, but the physical mechanisms underlying collective behavior in fibrous networks with strain-stiffening properties have not been investigated. Particularly intriguing is how local mechanical perturbations from contractile cells propagate through the nonlinear fibrous medium to coordinate long-range ordering, a process relevant to both physical models of active matter and biological processes such as wound healing and fibrosis.

A striking emergent behavior occurs in collagen matrices when cell spacing falls below a critical threshold: cells transition from minimal matrix remodeling to dramatic hydrogel compaction. Doha, et al.^5^ characterized this phase transition as a function of initial cell spacing, showing that cells spaced sufficiently close together coordinately compress the hydrogel to a small fraction of its initial volume, while cells beyond this critical distance fail to produce significant compaction. This phenomenon has been consistently observed across multiple systems with different cell types and collagen densities. Marquez, et al.^9^ found a critical cell density of *φ*_*c*_ = 400, 000 − 500, 000 cells/ml for chick embryo fibroblasts in 1 mg/ml type I collagen, corresponding to a spacing 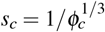 of 130 − 140 *µ*m. Similarly, Fernandez and Bausch observed a critical area density of 0.0001 − 0.0002 cells/*µ*m^2^ in 250 *µ*m thick gels (*φ*_*c*_ = 400, 000 − 800, 000 cells/ml) for MC3T3-E1 osteoblast cells in 2.4 mg/ml collagen, yielding *s*_*c*_ = 110 − 140 *µ*m. The most comprehensive characterization, by Doha, et al.^5^, demonstrated that contraction begins at 250,000 NIH 3T3 fibroblast cells/ml and increases to an asymptote at 2,000,000 cells/ml, establishing a critical spacing range of *s*_*c*_ = 79 − 160 *µ*m in 2 mg/ml collagen. The universality of this phenomenon extends beyond fibroblasts, as Xie, et al.^10^ showed that neuronal connectivity is similarly governed by intercellular spacing and collagen ECM mechanical properties.

This universality across different biological systems suggests underlying physical principles that transcend specific cellular details. The critical spacing threshold bears hallmarks of percolation phenomena in physics, where connectivity in disordered systems emerges abruptly at critical densities. In the case of cell-seeded matrices, the question becomes: what physical mechanism determines when cellular mechanical signals can percolate through the matrix to enable coordinated collective behavior?

To uncover the mechanisms driving these critical transitions, we developed an integrated bio-chemo-mechanical model that captures recursive interactions between cell contractility and ECM mechanics. Our approach combines two key components: first, a stress-responsive cellular model that accounts for mechano-chemical feedback driving myosin motor density and polarization^11–13^; and second, a fibrous ECM model that captures strain-dependent fiber recruitment, alignment, and stiffening, while accounting for the distinct responses of collagen to tension (stiffening) versus compression (buckling)^14–16^.

This integrated model reveals how cellular spacing determines the formation of mechanical tension bands between cells, establishing a threshold distance below which cells can effectively polarize their neighbors. Our simulations demonstrate that these cell-cell interactions depend crucially on the critical stretch ratio at which collagen fibers transition from compliant to strain-stiffening behavior—this parameter determines whether neighboring cells can establish sufficient mechanical cross-talk to coordinate tissue remodeling. By systematically analyzing these mechanical determinants, we provide a quantitative physical model for understanding how nonlinear elasticity controls phase transitions in active biological materials, with implications for both tissue mechanics and the broader physics of fiber-reinforced composite materials under active stress. Results also identify potential targets for therapeutic intervention in fibrotic disorders through manipulation of ECM mechanics.

## Fibroblasts can activate neighbors mechanically within a critical spacing

To identify how recursive cell-ECM interactions give rise to cell-cell mechanical communication, we combined a bio-chemo-mechanical model of fibroblasts with a nonlinear, strain-stiffening model of a fibrous ECM, and applied a computational implementation of this to model interactions between two ellipsoidal cells spaced a distance *s* apart axially within a much larger volume of ECM (Figure 1). The minor axes of the cells were *R* = 10 *µ*m. The contractile actomyosin machinery of the cells was modeled with a contractility tensor, ***ρ***, that evolved in response to the cell’s mechanical microenvironment^11,12^(see *Methods and materials*). This evolution accounted for chemo-mechanical feedback through a “chemo-mechanical feedback parameter,” *α*, representing the tendency to increase contractility in response to mechanical stress. It also accounted for the associated resistance to this increase in metabolic output, represented by the “chemical stiffness parameter,” *β* (see *Methods and materials*). The ECM was represented by the fibrous continuum constitutive model of Wang, et al.^17^, wherein the fibrous ECM is characterized by an elastic modulus *E*^*b*^ at low strain, and by transition to a strain-stiffening network for principal stretches beyond a critical value, *λ*_*c*_.

**Figure 1.**
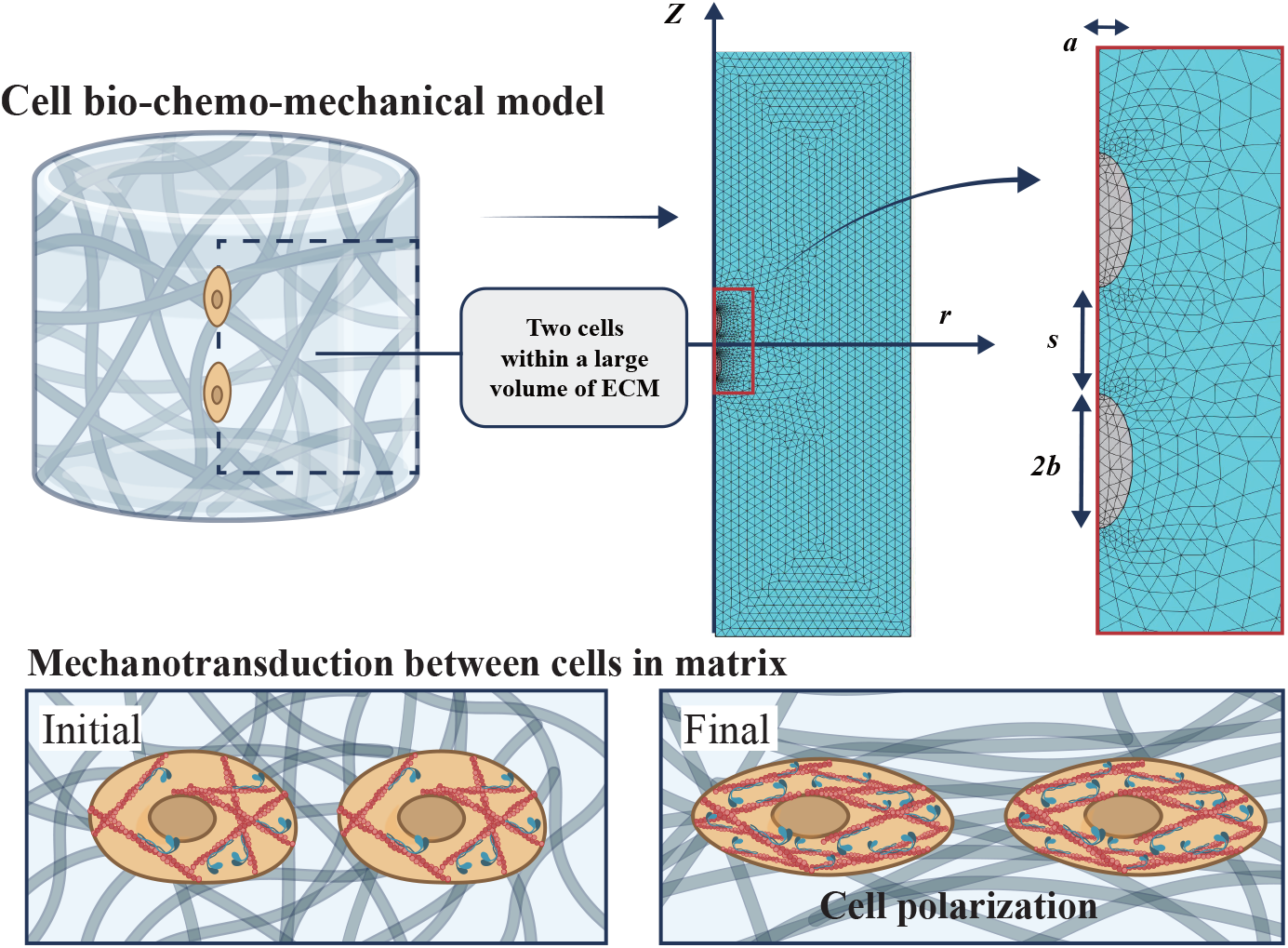
Bio-chemo-mechanical model of cell-ECM interactions and mechanical communication. (Top) Computational domain showing a pair of ellipsoidal cells embedded within a large fibrous extracellular matrix (ECM). The finite element mesh is highly refined near the cells to capture local mechanical gradients and fiber reorganization. (Bottom) Schematic of the integrated model incorporating: (i) a cellular contractility component that accounts for stress-dependent myosin motor synthesis and polarization, and (ii) a fibrous ECM model that captures strain-dependent fiber recruitment, alignment, and stiffening behaviors. This coupled framework enables prediction of how recursive interactions between cellular contractility and ECM remodeling establish the critical spacing threshold for mechanical communication.

Results shed light on ways that cells can become activated through mechanical interactions within their neighbors. As expected from numerous earlier studies^4,5,10,18–20^, even spherical cells in sufficiently close proximity stretched and aligned the collagen between them (Figure 2d-e). This was evident as bands of highly aligned collagen, with fibers aligning along the contractile axis between cells, and buckling perpendicular to this, as evident from the ratio between the maximum and minimum principal stretch ratios (*λ*_1_*/λ*_3_) (Figure 2e)^15,21^. The spherical cells compacted spherical regions of ECM around them to levels beyond the critical stretch, taken here to be *λ*_*c*_ = 1.04, as is appropriate for collagen in the 1-2 mg/ml range^15,17^. For cells spaced apart a distance *s* = 20*R*, these regions of strain-stiffening collagen did not overlap, but for cells spaced more closely together, the collagen matrix aligned strongly in the characteristic tension bands that are broadly reported^4,5,10,15,19^ (Figure 2d).

**Figure 2.**
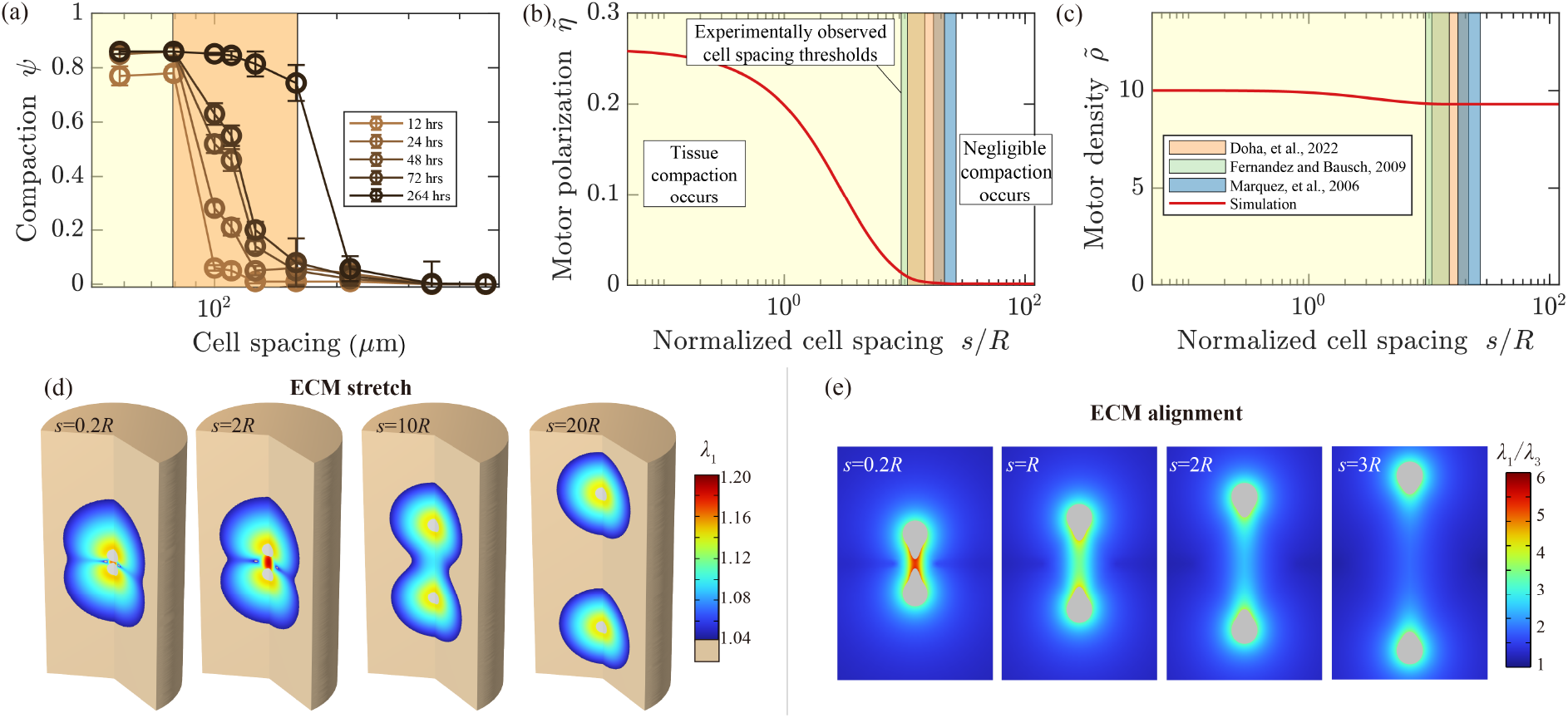
Critical spacing threshold governs collective cell behavior and tissue condensation. (a) Experimental data from Doha, et al.^5^ revealing a sharp phase transition in tissue compaction (*ψ*(*t*) = 1 − *A*(*t*)*/A*(*t* = 0), where *A*(*t*) is construct area) when cell spacing falls below the critical threshold of 79-160 *µ*m. (b-c) Our model predictions (red lines) accurately capture this threshold behavior observed across multiple independent studies (Supplemental Figure S1), with cell spacing normalized by nominal radius (10 *µ*m). The transition corresponds precisely to the distance at which myosin motor polarization 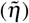 dramatically increases, while motor density 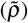 remains relatively constant, identifying polarization as the critical parameter governing collective behavior. (d) Mechanistic basis: cells below critical spacing generate overlapping zones of strain-stiffened ECM (where stretch exceeds the critical ratio *λ*_*c*_ = 1.04), creating tension bands that enable mechanical communication between neighbors. (e) Quantification of ECM fiber alignment (ratio of principal stretches *λ*_1_*/λ*_3_) shows dramatic reduction with increased cell spacing, demonstrating the loss of coordinated mechanical signaling beyond the critical distance. This reveals how discrete cellular interactions give rise to emergent tissue-scale behavior through mechanobiological feedback. Parameter values: *E* ^*f*^ */E*^*b*^ = 50, *λ*_*c*_ = 1.04, *α* = 2.4 kPa^−1^, *β* = 2.5 kPa^−1^, *ρ*_0_ = 1 kPa, *K* = 0.833 kPa, *µ* = 0.385 kPa^12^.

These effects were recursive, with the alignment and stiffening of the ECM leading to increasing activation of contractility in cells, and this contractility leading to further alignment of the ECM.We quantified these effects by considering two parameters, and analyzed them in the context of the extreme compaction of collagen when cells spacing is below the critical threshold (Figure 2b-c). The density of myosin motors, 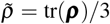, increased very slightly below the critical spacing (Figure 2c). However, the key difference was in the polarization of myosin motors within the cell (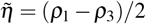, where *ρ*_1_ and *ρ*_3_ are the largest and smallest eigenvalues of ***ρ***, respectively. This changed dramatically at the critical spacing (Figure 2b). We therefore studied how cell-cell mechanical communication affected 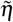 and 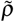.

### Myosin activation is determined by ECM properties alone, while cell polarization also depends upon cell spacing

To elucidate the mechanisms underlying these observations, we studied the joint effects on paratensile signaling of the cell spacing, *s/R*, and the pre-stiffening matrix modulus, *E*^*b*^. We again studied spherical cells so as to capture the first stages of cell activation. As *E*^*b*^ increased, the zone of strain stiffening around the cells decreased and the tension band between them disappeared (Figure 3a). However, despite this diminishing alignment of ECM fibers, cell activation (motor density, 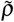) increased monotonically and substantially with *E*^*b*^ (Figure 3b). The effect of cell spacing on 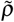 was only a few percent, and extremely small compared to the variation with respect to *E*^*b*^ (Figure 3d).

**Figure 3.**
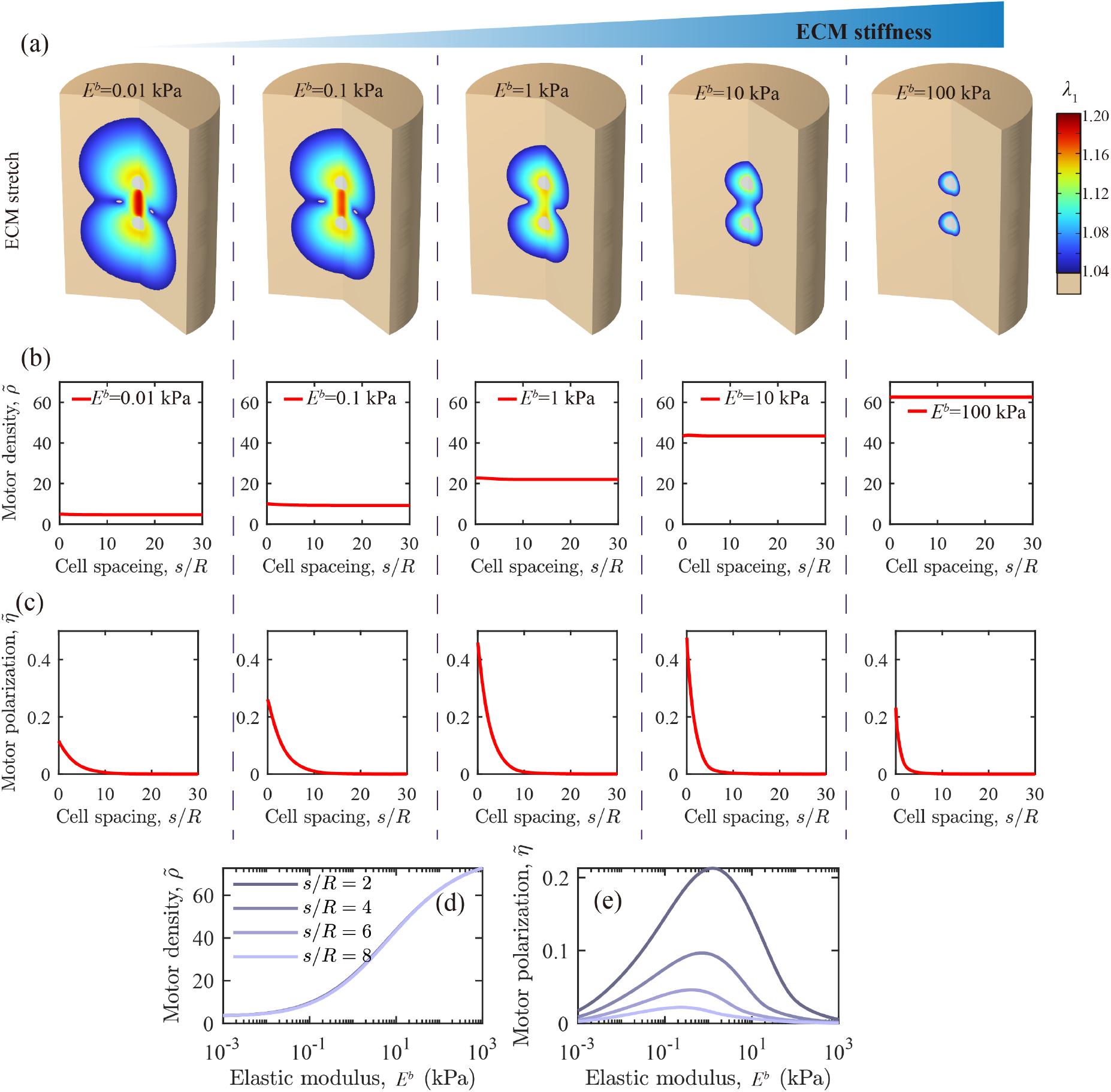
Competing mechanical effects create optimal stiffness for cell-cell communication. (a) Visualization of strain-stiffening regions and tension bands between cells at varying baseline matrix moduli (*E*^*b*^), revealing how increasing stiffness simultaneously shrinks the zone of mechanical influence while intensifying local activation (cell spacing *s/R* = 4). (b) Myosin motor density 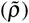 increases monotonically with matrix stiffness (*E*^*b*^) regardless of cell spacing, indicating that individual cell activation depends primarily on local matrix properties. (c) In contrast, myosin motor polarization 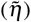 shows non-monotonic dependence on *E*^*b*^ and diminishes sharply beyond a critical spacing threshold that decreases with increasing stiffness, demonstrating the spatial limit of mechanical communication. (d) Quantitative comparison showing that motor density is almost exclusively determined by matrix stiffness while virtually independent of cell spacing, highlighting the distinct mechanisms governing activation versus polarization. (e) Peak polarization analysis reveals an optimal matrix stiffness that maximizes cell-cell communication, resulting from competition between enhanced cellular activation and diminished mechanical crosstalk with increasing stiffness. This competition mechanism explains why tissue condensation requires both sufficient cell density and appropriate matrix properties. Parameter values: *E* ^*f*^ */E*^*b*^ = 50, *λ*_*c*_ = 1.04, *α* = 2.4 kPa^−1^, *β* = 2.5 kPa^−1^, *ρ*_0_ = 1 kPa, *K* = 0.833 kPa, *µ* = 0.385 kPa^12^.

The effect of stiffness on myosin motor polarization 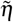 was much more substantial, with polarization disappearing beyond a critical spacing that diminished with increasing stiffness *E*^*b*^ (Figure 3c). The plateau signified the critical distance for mechanical communication between cells. The peak value of myosin motor polarization 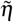 was a non-monotonic function of *E*^*b*^. This maximum was determined by a trade-off between having a modulus sufficiently low for overlapping regions of ECM alignment to connect the cells, but sufficiently high to have substantial myosin motor activation, 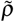 (Figure 3e). This balance thus determines how optimal stiffness arises through the effects on paratensile of cell spacing.

### Disruption of a mechanical feedback loop can halt fibrosis progression

To understand how these mechanical properties influence cellular behavior, we modeled a two-cell system within a large collagen matrix. Simulations demonstrated that both baseline Young’s modulus *E*^*b*^ and critical principal stretch *λ*_*c*_ affect cell polarization (Fig. 4a), a key indicator of cellular activation. Notably, our results suggested that merely modifying ECM stiffness might be insufficient to weaken cell-cell interactions, with decreases in *E*^*b*^ sometimes exacerbating polarization of fibroblasts for fibrotic tissue with moduli on the order tens of kPa^22^. These trends were robust against the parameters of the integrated cell-ECM mechanobiological feedback model (Supplemental Figures S2-S3), and against cell spacing (Supplemental Figure S4). This might provide insight into why efforts to reduce ECM stiffness using approaches such as upregulation of MMPs have not proven successful^23^.

**Figure 4.**
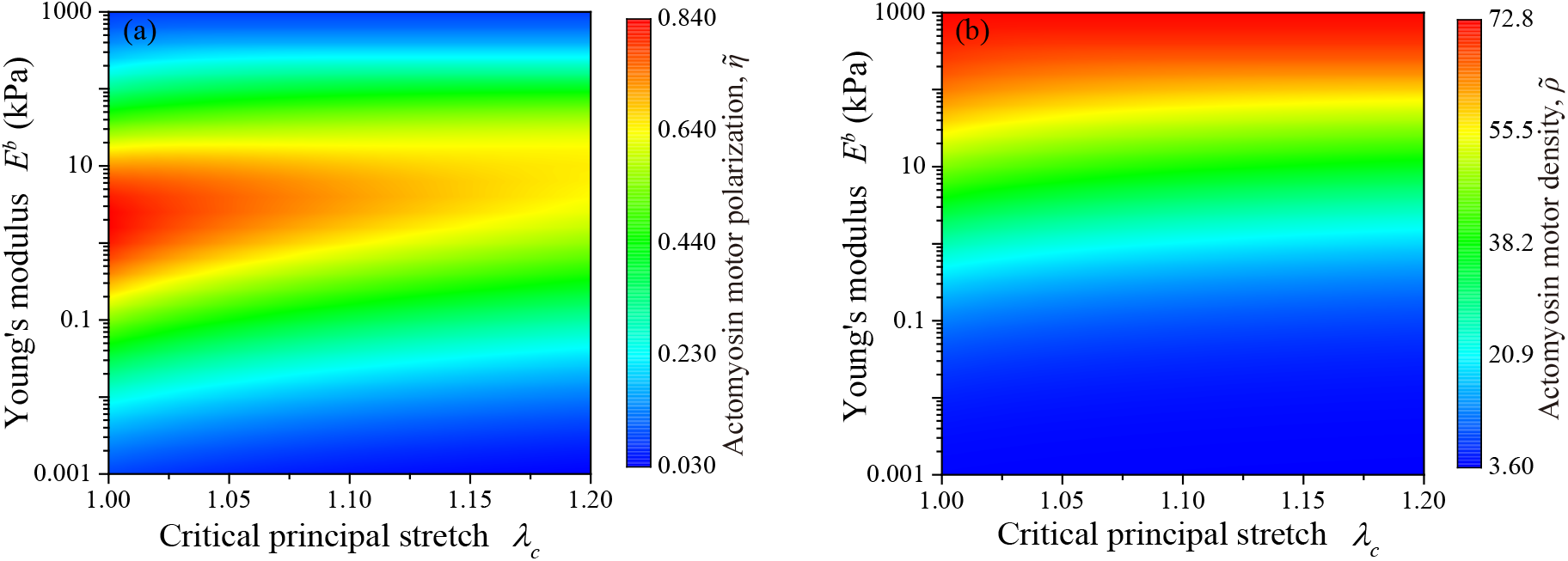
Joint effects of matrix properties on cellular mechanosensing. Effects of extracellular matrix (ECM) properties on (a) actomyosin motor polarization 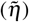 and (b) myosin motor density 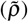 across a range of baseline moduli (*E*^*b*^) and critical stretch ratios (*λ*_*c*_). The heat maps reveal a complex, non-monotonic relationship between matrix mechanical properties and cell activation states. Notably, simultaneous modification of both stiffness and strain-stiffening behavior produces more dramatic changes in cell polarization than altering either parameter alone, suggesting that the mechanical feedback loop driving fibrosis progression depends on multiple ECM characteristics. This provides insight into why interventions targeting single mechanical aspects may show limited efficacy. Parameter values: *E* ^*f*^ */E*^*b*^ = 50, *α* = 2.4 kPa^−1^, *β* = 2.5 kPa^−1^, *ρ*_0_ = 1 kPa, *K* = 0.833 kPa, *µ* = 0.385 kPa^12^.

However, reducing crosslink density simultaneously decreased matrix elasticity and increased *λ*_*c*_, making the matrix more resistant to strain-stiffening. This combined effect led to reduced cell activation (Fig. 4b), affirming that targeting ECM crosslinkers could be an effective strategy for interrupting the mechanobiological feedback loop driving fibrosis progression.

### Cell spreading enhances mechanical feedback between cells and ECM

Because these results revealed a key role for actomyosin polarity, we next studied the polarization of shape that is characteristic of cell activation and spreading. An isolated cell with an aspect ratio of *b/a* = 2 in a very large ECM was modeled, with all parameters set at baseline level except the ECM modulus *E*^*b*^. Volume was kept constant so that *a*^2^*b* = *R*^3^. As with spherical cells, the zone around the cell for which the critical stretch ratio was exceeded shrank with increasing matrix modulus *E*^*b*^ (Fig. 5a). The displacement field was again a strong function of *E*^*b*^ (Fig. 5b). The myosin motor density 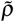 was again a strong function of *E*^*b*^, but only weakly dependent on *b/a* (Fig. 5c). Motor polarization 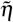 was a much stronger function of modulus, *E*^*b*^, with higher aspect ratio cells having higher motor polarization. The optimal stiffness for motor polarization decreased slightly with increasing aspect ratio (Fig. 5d).

**Figure 5.**
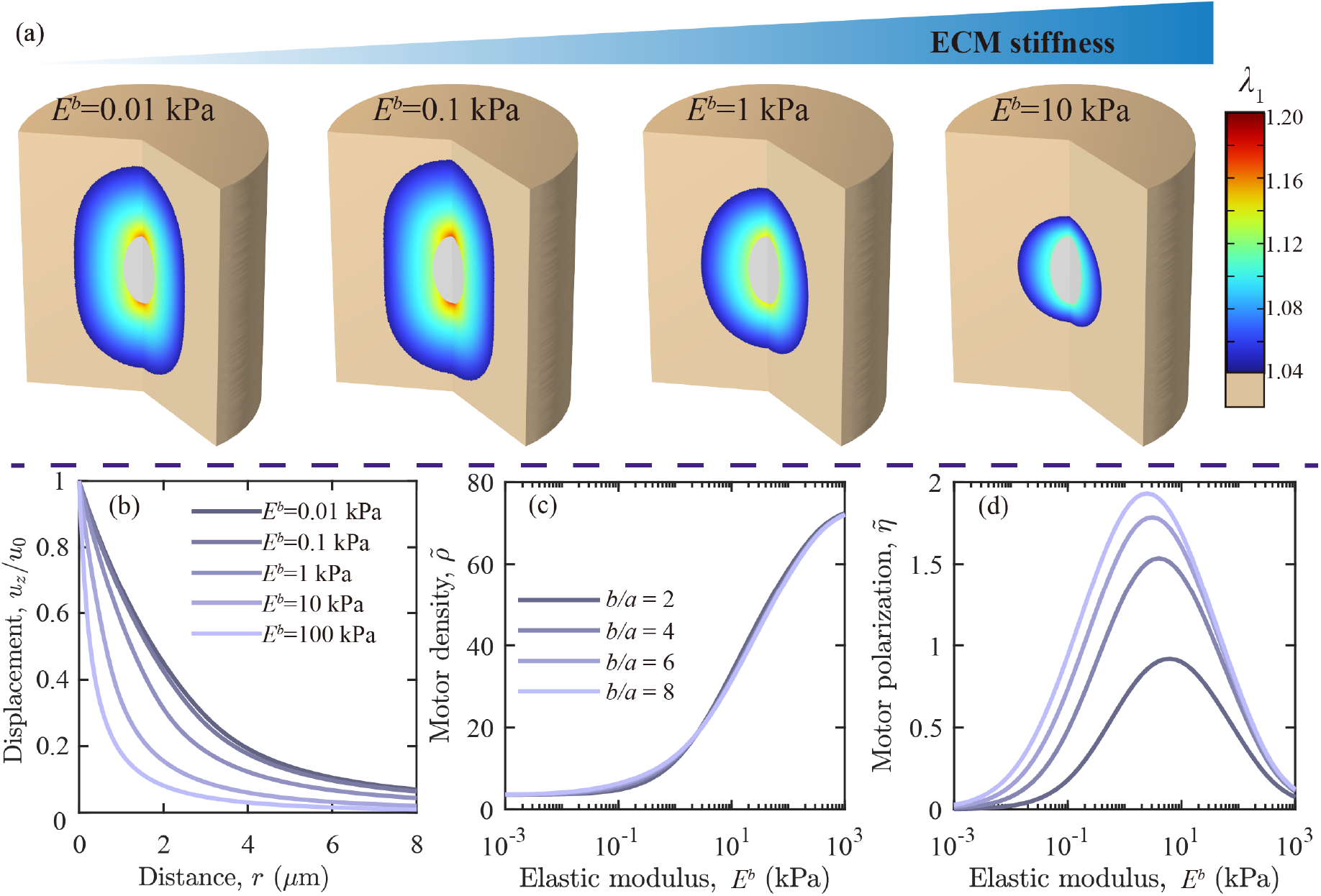
Cell geometry modulates mechanosensitivity to matrix stiffness. Analysis of how matrix rigidity affects single-cell mechanical behavior across different cellular aspect ratios. (a) Visualization of the critical stretch ratio zone around an elongated cell (*b/a* = 2) across a range of ECM moduli (*E*^*b*^), showing diminishing mechanical influence with increasing stiffness. (b) Spatial distribution of the displacement field generated by cell contractility, revealing how matrix stiffness constrains force transmission. (c) Myosin motor density 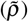 increases monotonically with *E*^*b*^ and exhibits modest enhancement with cellular elongation, demonstrating how cell shape amplifies mechanosensing. (d) Motor polarization 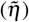 displays a non-monotonic response to matrix stiffness with a distinct optimum that shifts with cell aspect ratio, indicating that elongated cells achieve maximal polarization at lower matrix stiffness. This relationship explains why activated fibroblasts with higher aspect ratios can maintain polarization even as they stiffen their surrounding matrix. Parameter values: *E* ^*f*^ */E*^*b*^ = 50, *λ*_*c*_ = 1.04, *α* = 2.4 kPa^−1^, *β* = 2.5 kPa^−1^, *ρ*_0_ = 1 kPa, *K* = 0.833 kPa, *µ* = 0.385 kPa^12^.

### Cell spreading and elongation enhances force transmission and mechanical communication

Results suggest that that cell aspect ratio significantly modulates force transmission and cell-cell mechanical communication through the ECM. Increasing cell aspect ratio (*b/a*) while holding cell volume constant led to an expansion of the volume of ECM stretched beyond the critical stretch ratio (Fig. 6). This resulted in a characteristic dumbbell-shaped zone of influence around elongated cells, indicating enhanced ECM deformation.

**Figure 6.**
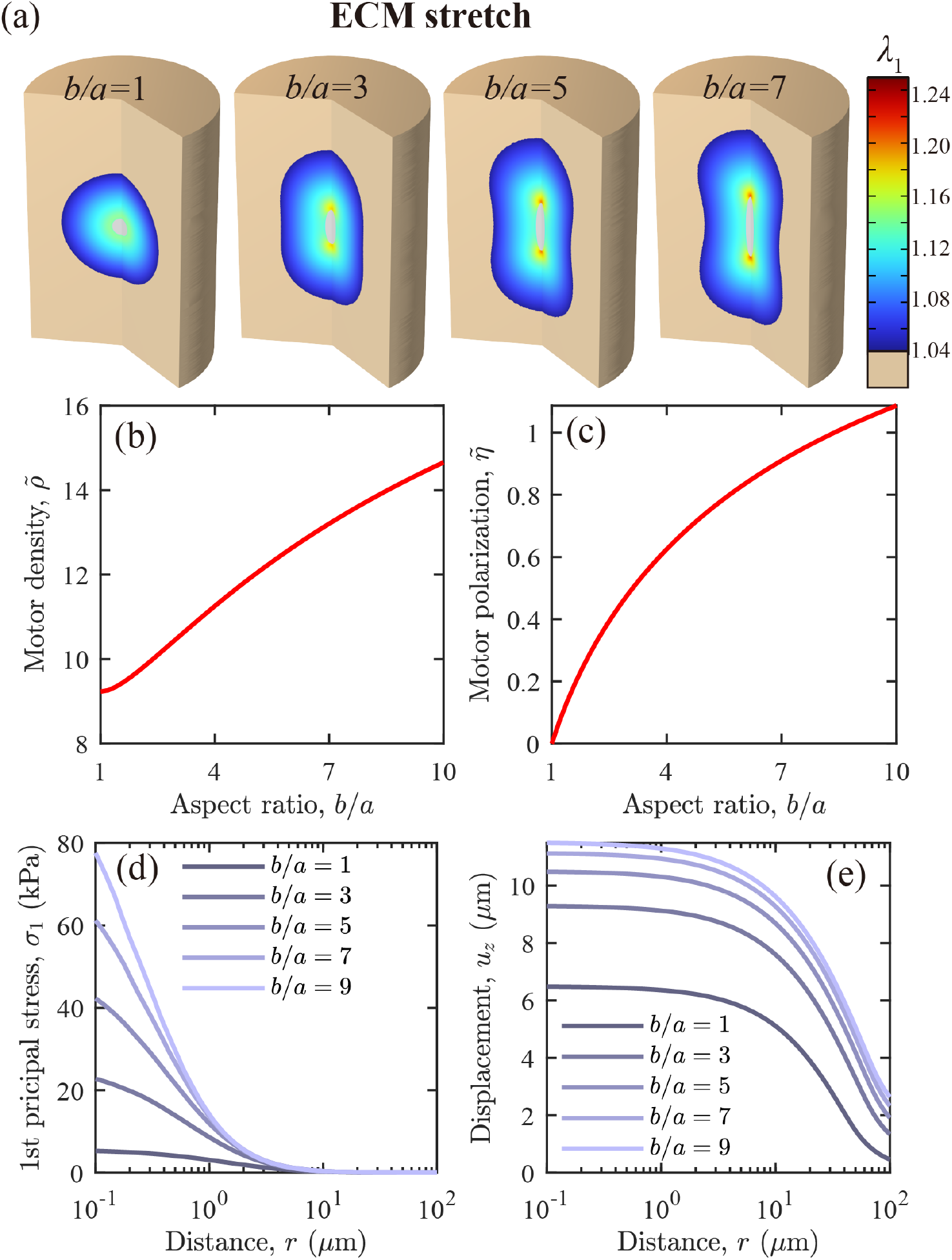
Cell elongation extends mechanical communication range. (a) Visualization of ECM regions exceeding the critical stretch ratio (*λ*_*c*_) for cells of increasing aspect ratio (*b/a*), revealing characteristic dumbbell-shaped zones of influence that extend significantly farther for elongated cells. (b-c) Quantitative analysis showing how cell elongation enhances both myosin motor density 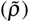 and polarization 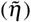, with polarization exhibiting particularly dramatic increases. (d-e) Spatial mapping of force transmission through (d) first principal stress and (e) displacement fields, demonstrating that elongated cells generate mechanical signals that propagate substantially farther through the matrix. These findings reveal a mechanism by which cell spreading and elongation—hallmarks of activated fibroblasts—effectively increase the range of cell-cell mechanical communication by amplifying long-distance force transmission. This geometric control of mechanosignaling range provides insight into how relatively sparse cells can coordinate tissue-scale responses during wound healing and fibrosis progression. Parameter values: *E*^*b*^ = 0.1 kPa, *E* ^*f*^ */E*^*b*^ = 50, *λ*_*c*_ = 1.04, *α* = 2.4 kPa^−1^, *β* = 2.5 kPa^−1^, *ρ*_0_ = 1 kPa, *K* = 0.833 kPa, *µ* = 0.385 kPa^12^.

Both mean motor density 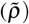 and mean motor polarization 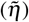 increased as cells became more elongated (Fig.6b-c). This enhancement in myosin motor activity coincided with higher curvature at the ends of elongated cells, which creates localized stress concentrations. These stress concentrations, in turn, triggered a bio-chemo-mechanical feedback loop within the cell, leading to increased overall contractility and motor density.

The model also demonstrated increased force transmission by elongated cells. Both the first principal stress and displacement were transmitted further by cells with higher aspect ratios (Fig. 6d-e), potentially facilitating enhanced cell-cell communication through the ECM.

The spatial anisotropy of stress in elongated cells led to increased expression of phosphorylated myosin light chain and F-actin at the cell ends. This localized upregulation resulted in higher motor polarization, further contributing to the cell’s exertion and transmission of forces. The strain-stiffening properties of the ECM amplified this effect, as the larger strains induced by elongated cells lead to increased local ECM stiffness.

## Discussion

Our results reveals a universal physical mechanism governing the abrupt phase transition in cell-seeded fibrous matrices. We demonstrate that a critical cell spacing threshold emerges from the competition between two opposing mechanical effects: matrix stiffness enhances individual cell activation while simultaneously attenuating long-range force transmission. This competition creates a mechanical bifurcation that manifests as dramatic tissue condensation when cells are spaced below approximately 80-160 *µ*m, in precise agreement with experimental observations across multiple independent studies^5,9,24^.

From the perspective of our model, the critical spacing threshold represents a mechanical percolation limit at which cell-generated tension bands become sufficiently interconnected to form a network capable of coordinating collective behavior. These tension bands arise when the matrix between neighboring cells is stretched beyond a critical stretch ratio (*λ*_*c*_), triggering strain-stiffening that provides a high-stiffness communication channel between cells^15,17^. Our bio-chemo-mechanical model quantitatively predicts how this critical spacing depends on matrix mechanical properties, particularly the critical stretch ratio that marks the transition from compliant to strain-stiffening behavior in fibrous networks^16,21^.

Cell elongation effectively extends this critical spacing by enhancing the spatial reach of mechanical signals. The stress concentrations at elongated cell tips generate localized regions of high strain that penetrate deeper into the surrounding matrix, effectively increasing the percolation radius of each cell. This mechanism explains why cellular processes that drive elongation, such as fibroblast activation, can actively modulate the range of mechanical communication and thereby control the phase boundary of the condensation transition^10,11^.

Our findings establish a quantitative framework for understanding how nonlinear material properties control emergent collective behavior in active biological systems. The competing effects of matrix stiffness on local activation versus long-range communication provides a physical explanation for the optimal stiffness observed in various biological processes^6–8^. This physical principle extends beyond biological systems to engineered active materials, where the interplay between structural nonlinearity and active force generation may similarly lead to emergent spatial ordering and abrupt phase transitions^4,12^.

The critical spacing threshold we identify represents a fundamental physical constraint on mechanical coordination in fibrous networks, with broad implications for understanding tissue development, pathological processes like fibrosis^4,20^, and the design of bio-inspired adaptive materials. By revealing how architectural and mechanical properties of fiber networks control phase transitions in active systems, our work bridges the gap between microscopic interactions and macroscopic collective behavior, a central challenge in both biological physics and materials science^1,2,5^.

## Acknowledgements

X.P. and X.-Q.F. acknowledge support from Natural Science Foundation of China (Grants 11921002, 12402364 and 12032014), as does Y.D. (Grants XXX). G.M.G. acknowledges support from the National Institutes of Health (R01AR077793) and the Human Frontier Science Program (HFSP-RGP016/2024).

## Author contributions

XP and YH performed simulations; all authors interpreted data; XP and GMG wrote the manuscript; all authors edited and approved the manuscript.

## Supplementary information

## Model of cell contractility and activation

Cells were modeled as actively contracting and adapting ellipsoids embedded in an initially random, fibrous ECM that was very large compared to the cell. Actomyosin contractility was modeled as a directionally-distributed field of myosin force dipoles that contract an isotropic background of actin filaments^11^. Following Shenoy, et al.^12^, this was represented by the contractility tensor, ***ρ***, derived by integrating forces from myosin dipoles over their orientation distribution. ***ρ*** is work conjugate to the linearized strain tensor, ***ε***, so that the specific work (per unit volume) done by phosphorylated myosin motors is *W* = ***ρ*** : ***ε***. Cellular contractility developed in response to tension exposure, resulting in increased F-actin polymerization and elevated levels of phosphorylated myosin light chain (p-MLC), as represented by evolution of ***ρ*** evolved with tension^11^. The enthalpic state of the cell was represented by five terms^12^:

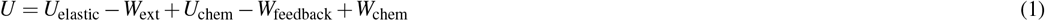

where the strain energy density is:

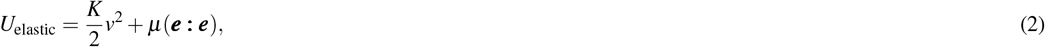

in which *K* is the cell’s bulk modulus, ***ε*** is the infinitesimal strain tensor, *v* = *tr*(***ε***) is the trace of ***ε***, *µ* is the shear modulus of the cell, and 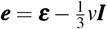 is the deviatoric portion of the strain tensor, in which ***I*** is the second order identity tensor. The external specific work was written:

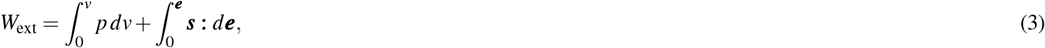

in which ***σ*** is the linearized stress tensor, 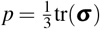, and ***s*** = ***σ*** − *p****I*** is the deviatoric component of the linearized stress tensor. The chemical energy density associated with phosphorylated myosin motors was written:

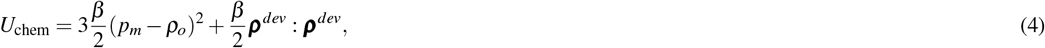

in which 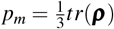, *ρ*_*o*_ is the baseline contractility, ***ρ***^*dev*^ = ***ρ*** − *p*_*m*_***I*** is the deviatoric component of ***ρ***, and *β* is a constant termed the “chemical stiffness.” Cells respond to mechanical stress by increasing ***ρ***, as governed by the feedback-associated specific work:

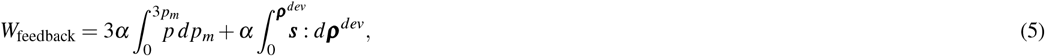

in which the constant *α* is the chemo-mechanical feedback parameter. Finally, the specific work done by the myosin motors was:

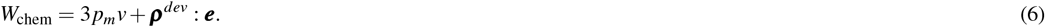

The temporal evolution equations can be inferred by taking time derivatives of Eq. 1^12^. For the cases of interest in which the timescales associated with motor recruitment and activation are short compared to the observation time, the following steady state conditions govern the active cell responses to mechanical stress:

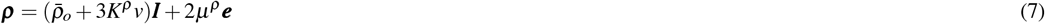

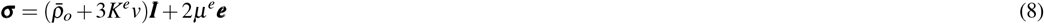

where 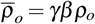, *K*^*e*^ = γ (*K* β − 1/3), *µ*^*e*^ = *γ*(*µβ* −1*/*2), *K*^*ρ*^ = *γ*(*Kα* −1*/*3), and *µ*^*ρ*^ = *γ*(*µα* −1*/*2), in which *γ* = 1*/*(*β* −*α*).

### Model of fibrous ECM mechanics

We modeled the fibrous ECM using the approach of Wang, et al.^17^, which treats the fibrous ECM as a superposition of elastic ground material and a set of fibers that provide additional resistance to deformation after stretching beyond a critical stretch level, *λ*_*c*_. The total strain energy density of the collagen network is *U* = *U*_*b*_ +*U*_*f*_, where the strain energy density associated with the ground material follows a neo-Hookean model:

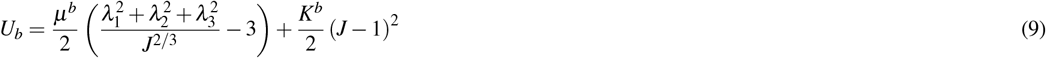

in which *µ*^*b*^ = *E*^*b*^*/*(2(1 + *ν*^*b*^)) is the initial shear modulus, *K*^*b*^ = *E*^*b*^*/*(3(1 − 2*ν*^*b*^)) is the initial bulk modulus, and *E*^*b*^ and *ν*^*b*^ are the initial elastic modulus and Poisson ratio of the ECM, respectively; {*λ*_1_, *λ*_2_, *λ*_3_} are the principal stretches; and *J* = *λ*_1_*λ*_2_*λ*_3_. The strain energy density associated with aligned fibers is:

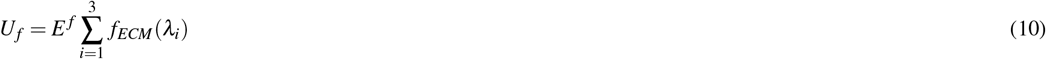

where *E* ^*f*^ is a strain-stiffening modulus, and the dimensionless strain-stiffening function *f*_*ECM*_(*λ*_*i*_) captures two fundamental principles: (*i*) the ECM exhibits strain stiffening solely in the direction of the first principal stretch when the stretch ratio exceeds a critical value *λ*_*c*_, and (*ii*) effects of strain stiffening are negligible for stretch less than *λ*_*c*_. From this, the stress in the matrix can be written as the sum of the stress in the ground material and the stress in the fiber network, ***σ*** ^*ECM*^ = ***σ*** ^*b*^ + ***σ*** ^*f*^, where^17^:

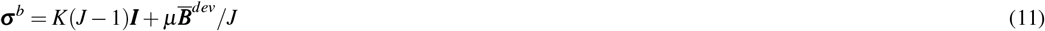

in which 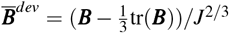 is the deviatoric part of the left modified Cauchy-Green tensor^17^. Here, 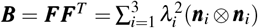, where ***F*** is the deformation gradient tensor, *λ*_*i*_ (*i* = 1, 2, 3) are the principal stretch ratios, and principal directions ***n***_*i*_ are the unit eigenvectors of ***B***. Following ^17, 21^, the contribution to fiber stiffening was written:

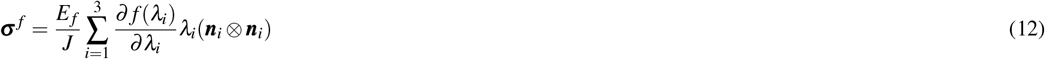

where:

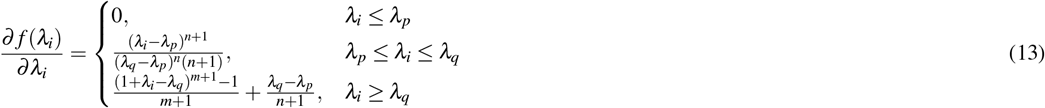

in which *λ*_*p*_ ≡ *λ*_*c*_ − *λ*_*t*_*/*2, *λ*_*q*_ ≡ *λ*_*c*_ + *λ*_*t*_*/*2, *λ*_*c*_ is the stretch ratio at which fiber stiffening begins to contribute strongly to ECM mechanics, and *λ*_*t*_ is a transition region. For and strain-stiffening exponents *m* and *n* are fitting parameters^15,21^.

### Numerical methods

Equations were solved using the finite element in the COMSOL Multiphysics (COMSOL, Inc., Burlington, MA) environment. Cells were treated as axisymmetric. Cells and ECM were discretized with linear, triangular elements. Mesh convergence studies were performed, and convergence was typically achieved in models with approximately 100000 elements.

COMSOL’s Solid Mechanics Interface (a component of COMSOL’s Structural Mechanics Module) was adopted to model the cross-talk between cells and ECM. Cell responses included passive and active components. The passive component was modeled as linear elastic. The active contractile stress, ***ρ***, was superimposed on the cell. To capture the presence of the two distinct families of aligned and isotropic fibers, a neo-Hookean model was adopted to capture the isotropic response, while Eq. 12 was used to include the contribution due to the aligned fibers.

## Supplemental Figures

**Figure S1.**
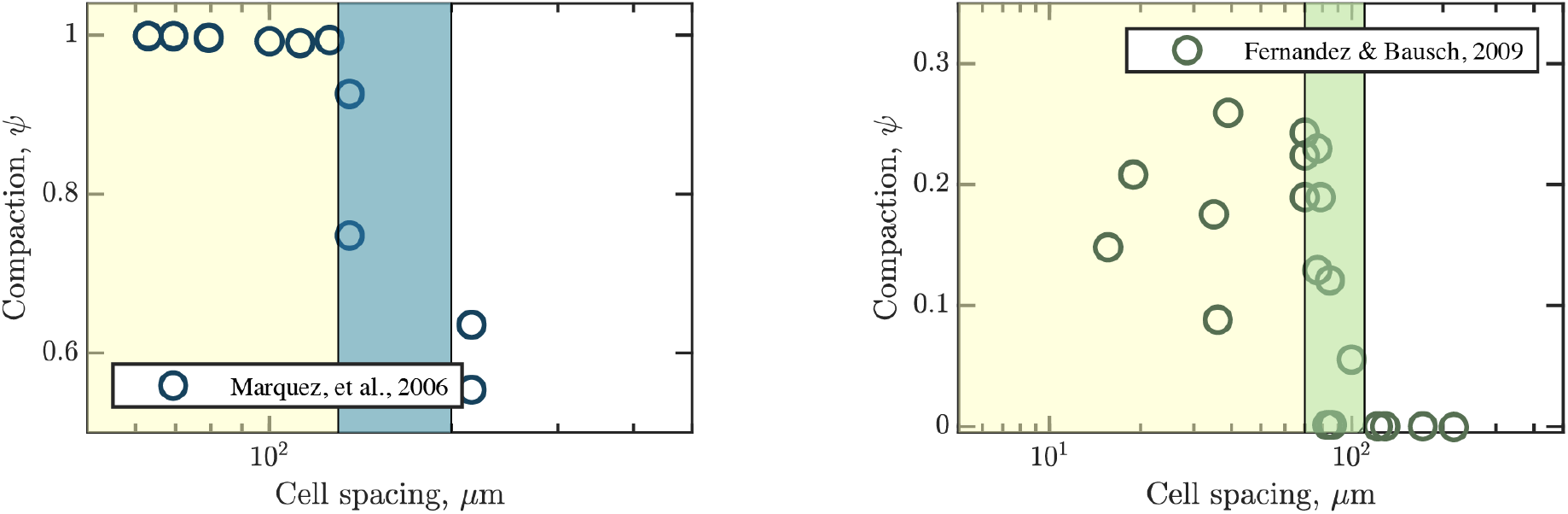
Experimental validation of critical cell spacing threshold across multiple cell types and collagen densities. (Left) Data from Marquez, et al.^9^ showing a sharp transition in tissue compaction at a critical cell density of *φ*_*c*_ = 400, 000 − 500, 000 cells/ml for chick embryo fibroblasts in 1 mg/ml type I collagen, corresponding to a critical spacing 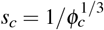 of 130-140 *µ*m. (Right) Independent confirmation from Fernandez and Bausch^24^ demonstrating a similar phase transition at a critical area density of 0.0001-0.0002 cells/*µ*m^2^ in 250 *µ*m thick gels (*φ*_*c*_ = 400, 000 − 800, 000 cells/ml) for MC3T3-E1 osteoblast cells in 2.4 mg/ml collagen, yielding *s*_*c*_ = 110 − 140 *µ*m. The consistency of this critical spacing range across different experimental systems supports the universal nature of the mechanical threshold governing tissue condensation.

**Figure S2.**
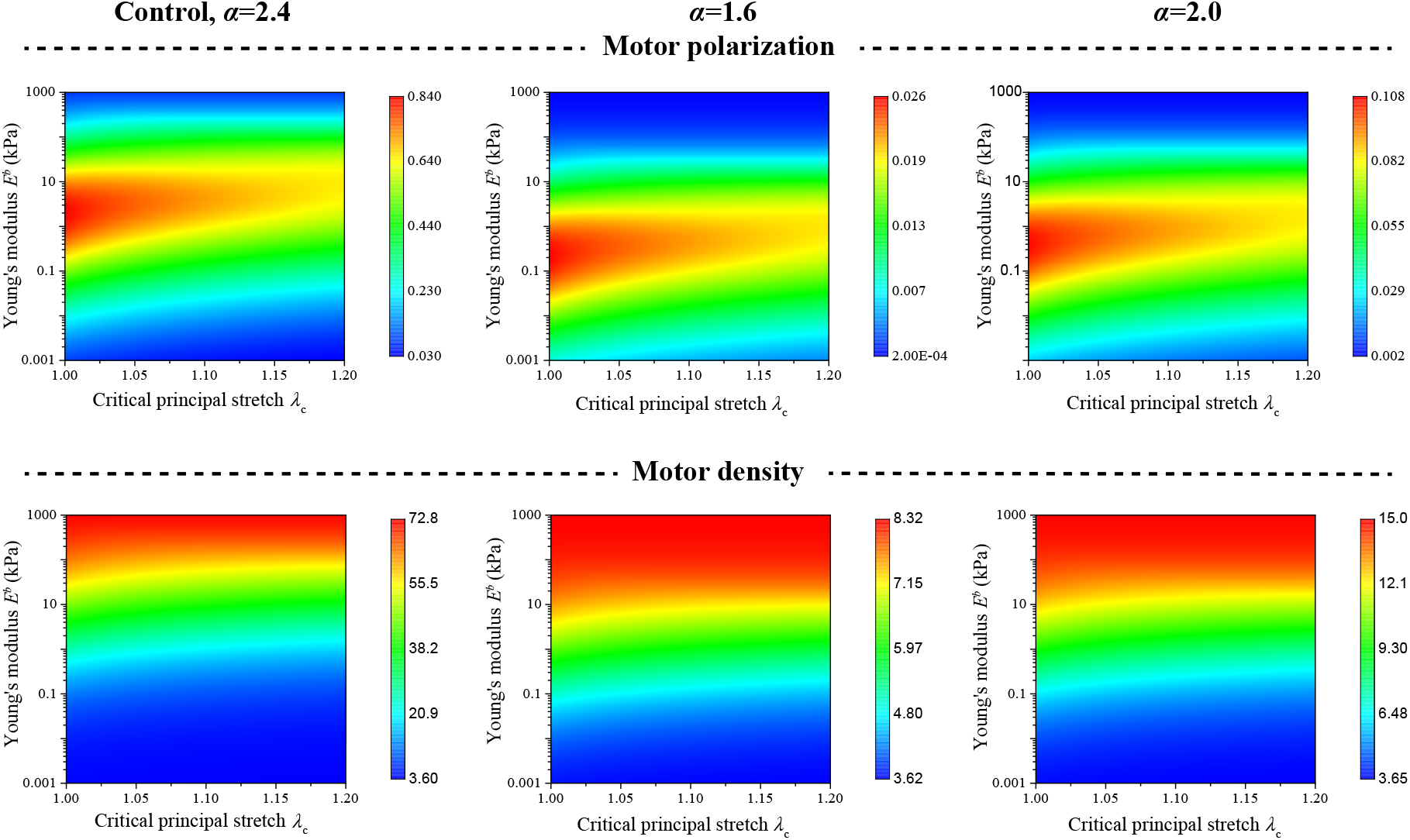
Model predictions are robust to variations in chemomechanical coupling strength. Sensitivity analysis of the computational model’s response to variations in the chemomechanical coupling parameter *α*, which governs cellular contractile response to mechanical stress. The predicted relationships between ECM properties (baseline Young’s modulus *E*^*b*^ and critical principal stretch *λ*_*c*_) and cell polarization remain qualitatively consistent across a range of *α* values. While motor density magnitudes vary with *α*, the fundamental trends persist, confirming that the model’s predictions about matrix mechanical effects are robust to uncertainty in this cellular parameter.

**Figure S3.**
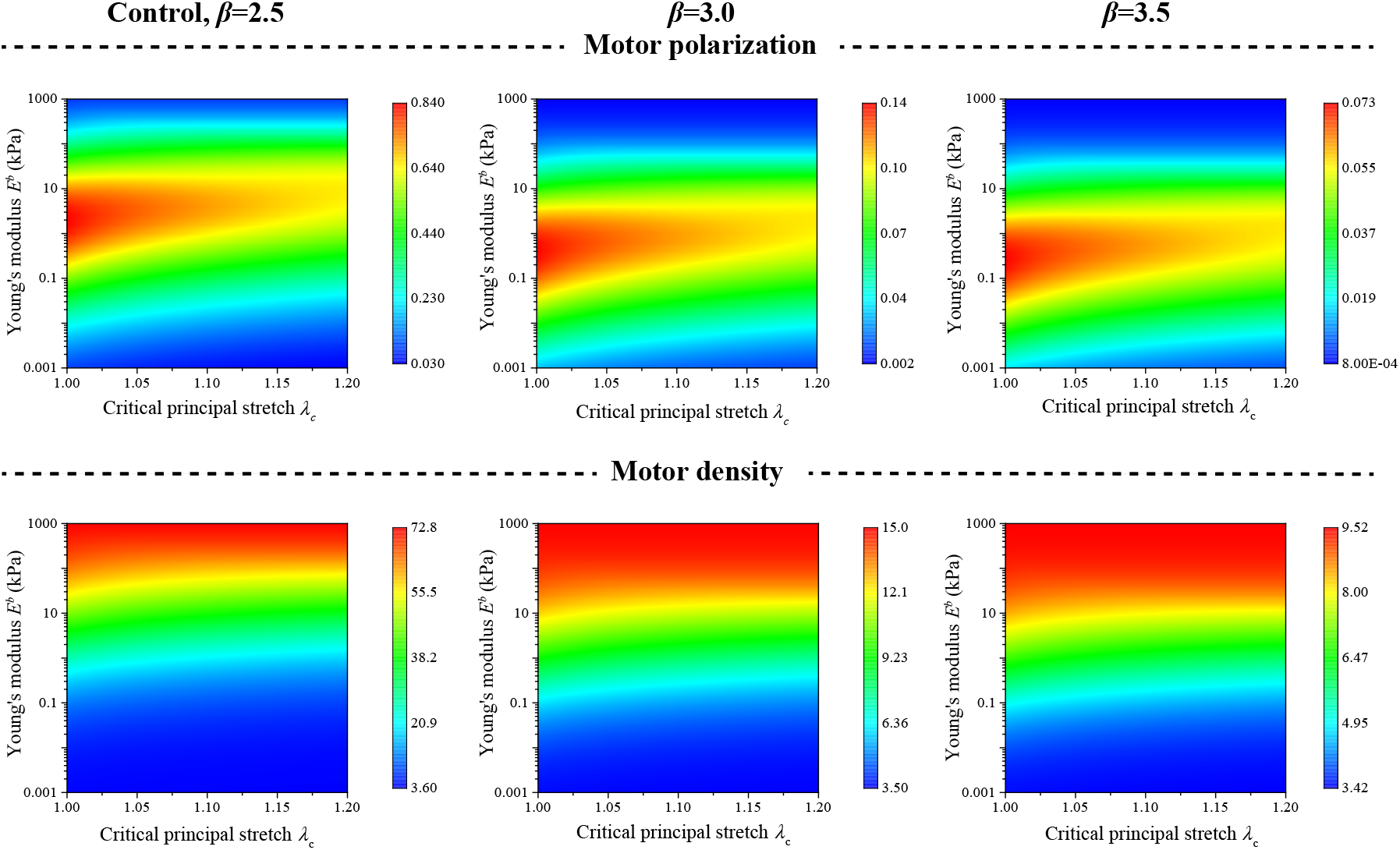
Model predictions are maintained across variations in cell polarization sensitivity. Sensitivity analysis of computational predictions to changes in the polarization parameter *β*, which determines how strongly cells align their contractile machinery with principal stress directions. While the optimal Young’s modulus (*E*^*b*^) for cell activation shifts with *β*, the qualitative relationships between ECM properties (*E*^*b*^ and critical principal stretch *λ*_*c*_) and cell polarization remain consistent. Motor density magnitudes vary with *β* but maintain similar trends, demonstrating that model predictions about matrix mechanical properties are robust to uncertainty in cellular mechanosensitivity.

**Figure S4.**
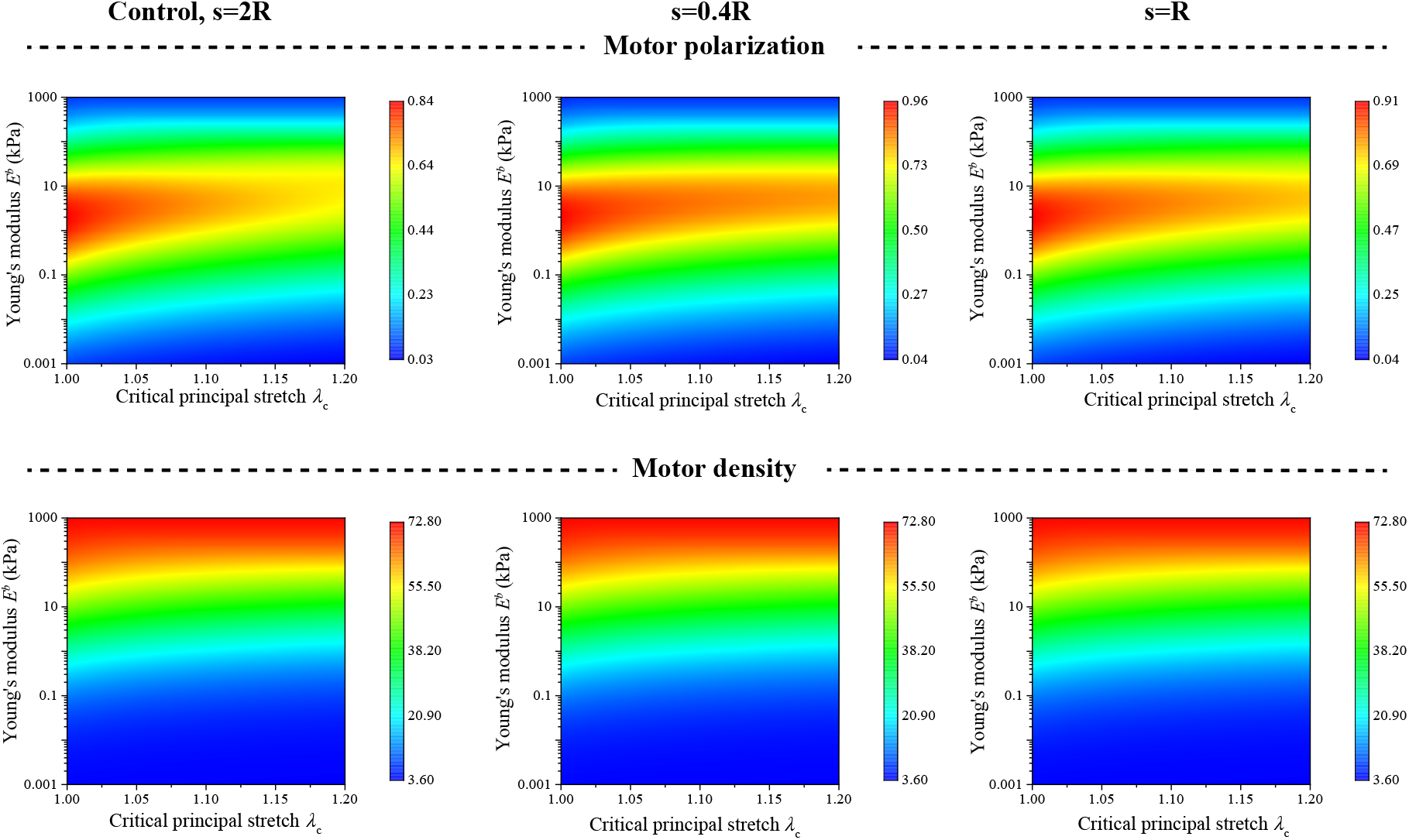
Model predictions are stable across different cell-cell separation distances. Sensitivity analysis of computational predictions to variations in cell-cell spacing (*s*), showing that key relationships between ECM properties (baseline Young’s modulus *E*^*b*^ and critical principal stretch *λ*_*c*_) and cell polarization persist despite changes in intercellular mechanical coupling. While the strength of cell-cell interactions naturally decreases with distance, the fundamental relationships between ECM properties and cell activation remain qualitatively unchanged, confirming that model predictions about matrix mechanical properties are robust to cellular spacing.

